## Supplemental Information for "Modelling how curved active proteins and shear flow pattern cellular shape and motility"

Shubhadeep Sadhukhan* and Nir Gov†
*Department of Chemical and Biological Physics, Weizmann Institute of Science, Rehovot, Israel*

Samo Penič‡
*Laboratory of Physics, Faculty of Electrical Engineering, University of Ljubljana, Ljubljana, Slovenia*

Aleš Iglič§
*Laboratory of Physics, Faculty of Electrical Engineering, University of Ljubljana, Ljubljana, Slovenia and Laboratory of Clinical Biophysics, Faculty of Medicine, University of Ljubljana, Ljubljana, Slovenia*
(Dated: March 26, 2023)

### S-1. SIMULATION DETAILS

The time evolution of the vesicle in our MC simulations consists of [2] (1) vertex movement, and (2) bond flip. In a vertex movement step, a vertex is chosen and attempts to move to a position randomly chosen within a sphere of radius $s$ drawn around the vertex position. We set $s = 0.15$ in units of $l_{\text{min}}$. In a bond flip movement, a bond is chosen common to two neighbouring triangles of the triangulated surface. Two such neighbouring triangles form a quadrilateral. The bond connecting two vertices in the diagonal direction of the quadrilateral is destroyed and recreated between the other two vertices, previously unconnected. We use the Metropolis algorithm to update the system—a statistical mechanics simulation method to study complex systems. The new state of the system increases the energy of the system by an amount $\Delta E$ is accepted with the rate $\exp(-\Delta E/k_BT)$. However, the new state is accepted certainly if such a move decreases the system energy.

In the simulations presented in this paper, we use the model parameters as follows: the total number of vertices $N = 1447$, the bending rigidity $\kappa = 20\ k_BT$, protein-protein interaction strength $w = 1\ k_BT$. The adhesion interaction length is set to be $l_{\text{min}}$. Among $N$ vertices, $N_c$ vertices are occupied by the curved membrane proteins. The spontaneous curvature at all the $N_c$ vertices is set to $c_0 = 1\ l_{\text{min}}^{-1}$. The spontaneous curvature for the rest of the vertices is set to zero. The percentage of curved protein vertices $\rho = 100N_c/N$ is an important parameter. Throughout this paper, we calculated the velocity of the center of mass in units of $l_{\text{min}}/2000$ MC steps whereas all the lengths are measured in units of $l_{\text{min}}$.

### S-2. MODELLING OF SHEAR FORCE

We modelled the force due to the shear flow such that the membrane feels a force that is tangential to the membrane surface that is perpendicular to the local normal of the membrane (Fig.1). The direction of the local shear force depends on the global shear flow direction $\hat{v}^{\text{shear}}$ and the local normal to the membrane $\hat{n}$. First, we find the direction which is perpendicular to both the flow direction and the local normal at the membrane by a cross product: $\hat{v}^{\text{shear}} \times \hat{n}$. The flow should pass normally through the plane formed by the local normal $\hat{n}$ and the direction $\hat{v}^{\text{shear}} \times \hat{n}$. Therefore, we modelled the direction of the force due to shear flow as: $\hat{n} \times (\hat{v}^{\text{shear}} \times \hat{n})$. We always set the strength of the shear force by the shear rate parameter $a$, and the distance to the adhesive surface $z_i - z_{\text{ad}}$. The magnitude of the force due to shear on a vertex positioned at $\overrightarrow{r_i} = (x_i,\ y_i,\ z_i)$ is $F^{\text{shear}} = a(z_i - z_{\text{ad}})$.

We simulated four cases with four different shear rates as shown in Fig S-1. Four columns correspond to a shear rate $a = 0,\ 0.005,\ 0.01,\ 0.012(k_BT/l_{\text{min}}^2)$. Four rows correspond to different cases with different parameter sets: (a) No proteins, (b) protein percentage $\rho = 3.45\%$ and no active force $F = 0$, (c) protein percentage $\rho = 3.45\%$ and active force $F = 0.5k_BT/l_{\text{min}}$, (d) protein percentage $\rho = 6.91\%$ and no active force $F = 0$. We kept the location of the centre of mass in the $x - y$ plane fixed for all the simulations discussed in this section.

*
†
‡
§

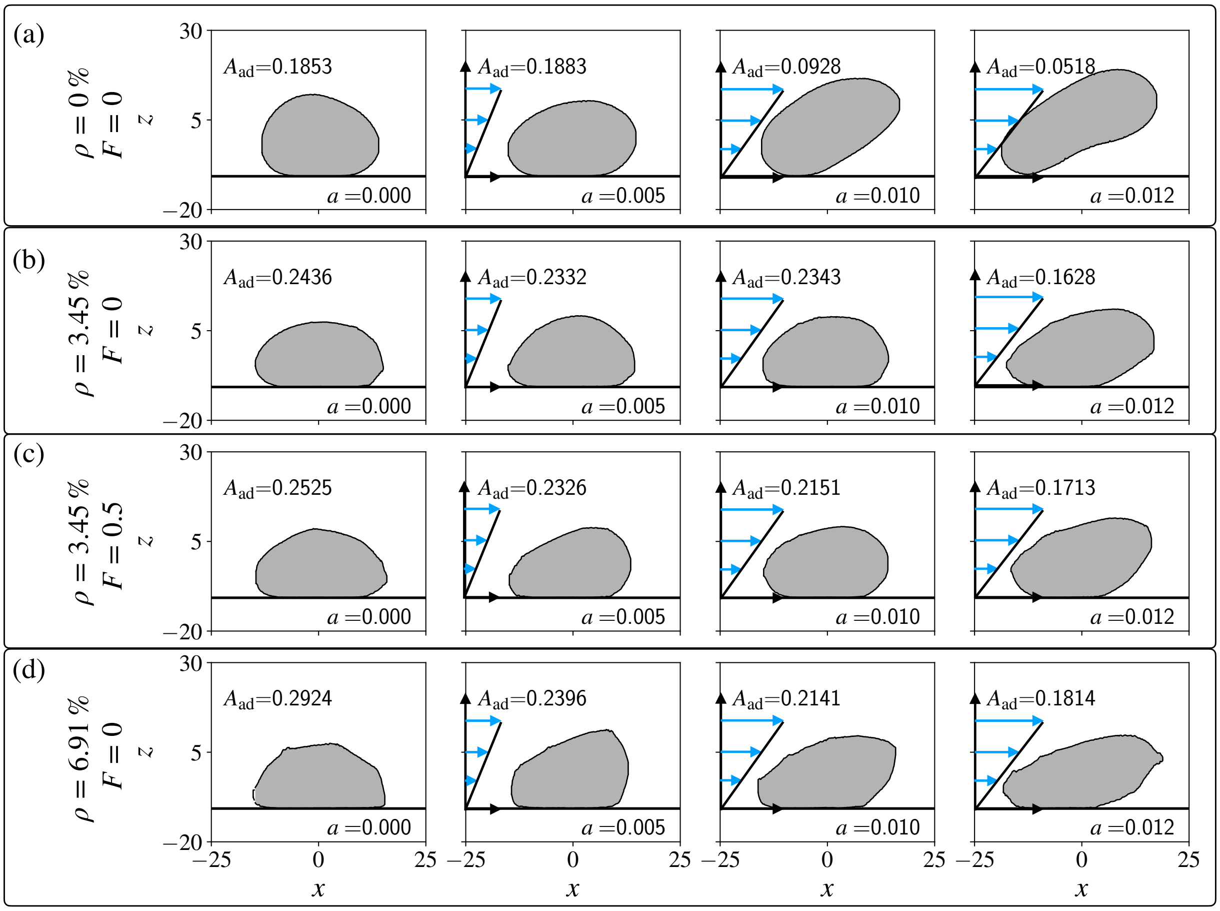


FIG. S-1. Cross-sectional area of the vesicle at the vertical plane that cuts through the center of mass at $y = y_{\rm avg}$, and we applied the shear force along the $\hat{x}$ direction. Each column corresponds to a different shear rate: $a = 0,\ 0.005,\ 0.01,\ 0.012(k_BT/l_{\rm min}^2)$. We set the adhesion strength $E_{\rm ad} = 0.5k_BT$. Four rows are corresponding to the parameter set (a) No protein case: $\rho = 0\%,\ F = 0$; (b) Passive proteins case: $\rho = 3.45\%,\ F = 0$; (c) Active proteins case: $\rho = 3.45\%,\ F = 0.5k_BT/l_{\rm min}$; (d) Passive proteins case with higher density: $\rho = 6.91\%,\ F = 0$.

We find how shear forces affect the shape of the vesicle by plotting the cross-section of the vesicles in the $x$-$z$ plane as shear flow is applied in the $x$ direction (Fig. S-1). The protein-free shapes that we obtain are very similar to the shapes found in [1], including the tendency of the vesicles to lift and eventually detach as the shear flow strength is increased. We calculated the magnitude of the adhered surface $A_{\rm ad}$ for each case shown in Fig. S-1 (given in each panel), which we find to be decreasing with the shear rate $a$. We find that the vesicle spreads more when the protein percentage $\rho$ is higher for any fixed shear rate. We also find that the vesicle with active curved proteins spreads more than the vesicle with the passive proteins [3], for a fixed protein percentage $\rho$ and the shear rate $a$.

### S-3. THE BODY AXIS

The body axis of an unstructured vesicle is defined as its most elongated internal axis, as follows. The vesicle has $N$ vertices. The $i$th vertex has the position vector $\overrightarrow{r_i} = (x_i,\ y_i,\ z_i)$. The centre of mass of the vesicle is given by the position vector $\overrightarrow{CM} = (CM_x,\ CM_y,\ CM_z)$. We find the matrix $G$ given by,

$$G = \begin{bmatrix} G_{xx} & G_{xy} \\ G_{yx} & G_{yy} \end{bmatrix}, \tag{S-1}$$

where, $G_{xx} = \sum_i (x_i - CM_x)^2, G_{xy} = G_{yx} = \sum_i (x_i - CM_x)(y_i - CM_y), G_{yy} = \sum_i (y_i - CM_y)^2$. Next, we found the eigenvector that corresponds to the largest eigenvalue, and it gives us the body axis along which the vesicle is elongated.

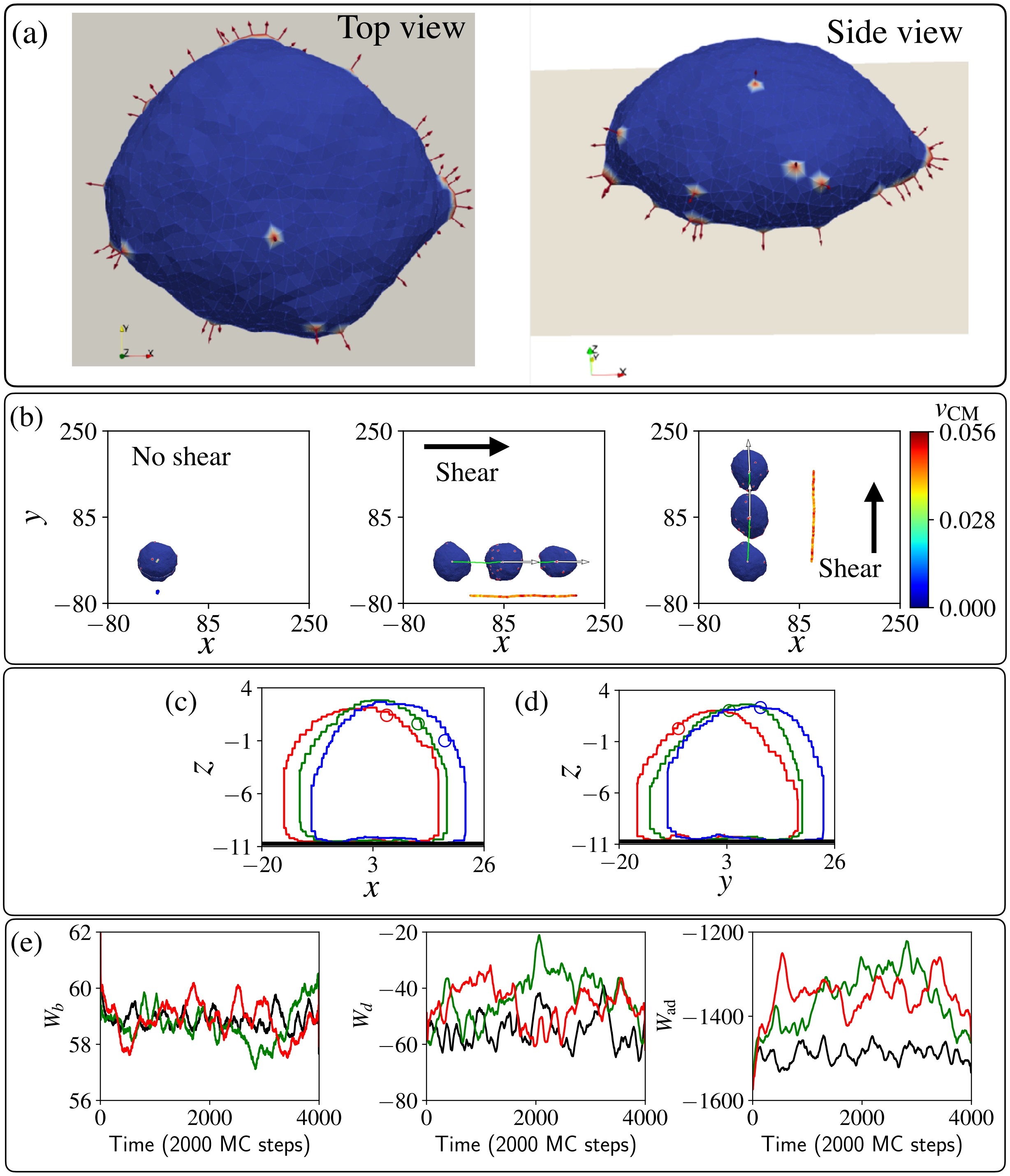


FIG. S-2. Nonpolar vesicle: (a) A weakly spread vesicle on a weakly adhesive surface and low active force [3] $F = 0.5k_BT/l_{\min}$, $E_{\rm ad} = 1k_BT, rho = 3.45\%$. (b) Trajectories of the vesicle for three different shear conditions (no shear, shear in $x$ direction, and shear in $y$ direction) in lime colour, in three respective columns. We set the shear rate $a = 0.01k_BT/l^2_{\min}$. A shifted trajectory is shown on each panel with a color code that shows the velocity of the centre of mass of the vesicle. (c) Cross-sections of the vesicle on $x$-$z$ plane when the shear is in the $x$ direction at Time =0, 75, and 100 (unit of 2000 MC steps) in red, green, and blue colours. A particular vertex is encircled on the cross sections. (d) Cross-sections of the vesicle on $y$-$z$ plane when the shear is in the $y$ direction at Time =0, 75, and 100 (unit of 2000 MC steps) in red, green, and blue colours. A particular vertex is encircled on the cross sections. (e) Time evolution of bending energy, protein-protein binding energy, and adhesion energy in three respective panels. Black, green and red lines correspond to the cases of no shear, shear in $x$ direction, and shear in $y$ direction respectively.

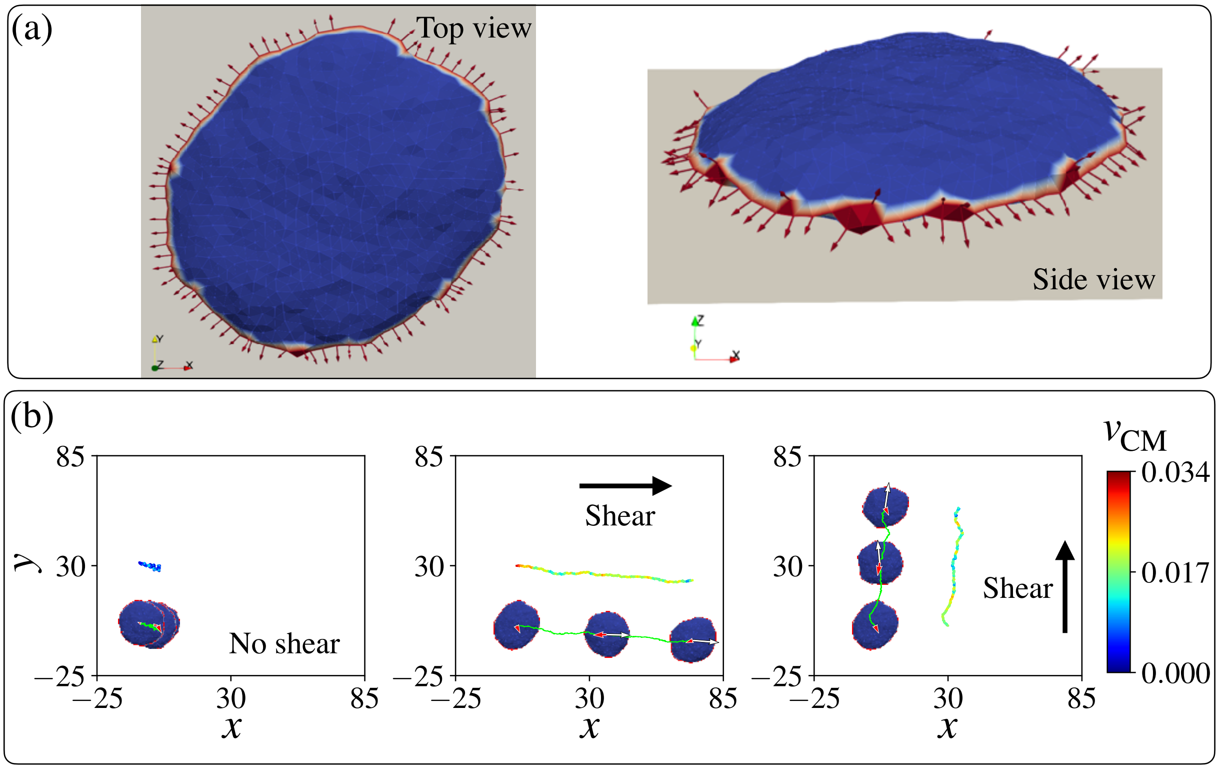


FIG. S-3. Pancake-shaped vesicle: (a) We set $F = 2k_BT/l_{\mathrm{min}}$, $E_{\mathrm{ad}} = 0.5k_BT$, $\rho = 6.91\%$ to get a pancake-shaped vesicle [3] (b) The trajectories (lime solid lines) of a pancake-shaped vesicle under different shear conditions in three different columns. We set the shear rate $a = 0.01k_BT/l_{\mathrm{min}}^2$ for the second and third panels. The active force and the total force are shown in red and white arrows respectively in each column. The velocity of the centre of mass is shown with a solid shifted trajectory with colours mapped to its velocity $v_{\mathrm{CM}}$ for each case.

### S-4. NON-POLAR SHAPED VESICLES UNDER SHEAR

There are several different kinds of non-motile vesicles that emerge in our model [3], other than the two-arc-shaped vesicle presented in the main text (Fig.3).

#### A. Weakly spread vesicle under shear

We can get non-motile non-spreading vesicles when the active force and adhesion strength are low ($F = 0.5\ k_BT/l_{\mathrm{min}}$, $E_{\mathrm{ad}} = 1k_BT$) and protein concentration is low $\rho = 3.45\%$ (Fig S-2a). The proteins are scattered randomly on the membrane, therefore they can not produce a significant and persistent force in a particular direction. Hence, the vesicle remains at the same position and fluctuates under thermal noise when no shear flow is present, as shown in the first column of Fig S-2(b). We set the shear rate $a = 0.01\ (k_BT/l_{\mathrm{min}}^2)$ for all the sheared cases in this section. The vesicle rolls with the flow as shown in Fig S-2(b), where shear is in the $x$ and $y$ direction respectively.

Next, we shed some light on the question of how the vesicle is moving with the shear flow. We plot the cross-section of the vesicles on the vertical plane through their center of mass, and the direction of shear flow in Figs. S-2(c,d). By following the motion of a particular vertex on the cross-section we see that the vesicle moves with the flow, performing a mixture of sliding and rolling.

Next, we plot the bending energy $W_b$, protein-protein binding energy $W_d$, and adhesion energy $W_{\mathrm{ad}}$ of the system in Fig S-2(e). It is very clear that adhesion is stronger in the absence of shear. It indicates the tendency of shear to cause a lifting force and de-adhesion of the vesicle (as we showed also in Fig.S-1) is the dominant effect.

#### B. Pancake-shaped vesicle under shear

If we increase the protein density in such a way that the proteins can form a closed circular cluster around the rim of the vesicle, we can get another type of non-motile vesicle, i.e. a pancake-like shape (Fig. S-3a). In order to get a pancake-shaped vesicle we set a higher active force strength $F = 2k_BT/l_{\rm min}$, adhesion strength $E_{\rm ad} = 0.5k_BT$ and most importantly a higher protein percentage $\rho = 6.91\%$.

As the active force is equal in every direction due to the formation of the circular rim cluster (Fig. S-3a), the net force on the vesicle is zero. Therefore, without any other force, this shape is non-motile on the adhesive substrate, as shown in Fig. S-3(b). However, the addition of the shear force breaks the force balance, and the vesicle slides with the flow. We studied such pancake-shaped vesicles under the shear flow along $x$ and $y$ directions respectively. The trajectories of the centre of mass of the vesicles under such shear flow show that the vesicle is moving with the flow as shown in Fig. S-3(b). We do not observe significant polarization of the vesicle with the flow, so that the internal active forces do not contribute to the shear-induced migration.

### S-5. MOVIES

**Movie 1: Polar vesicle** Polar vesicle is obtained using parameters $E_{\rm ad} = 3k_BT$, $F = 2k_BT\ l_{\rm min}^{-1}$, and $\rho = 3.45\%$. Four panels for four different shear conditions. When the shear flow is absent, the motile vesicle moves in the $x$ direction persistently. It shows the U-turn of the polar vesicle when $\hat{v}^{\rm shear} = \hat{x}$. Protein aggregate rotates and the vesicle moves against the shear when $\hat{v}^{\rm shear} = \hat{y}$.

**Movie 2: Two-arc vesicle** Non-motile two-arc-shaped vesicle is obtained $E_{\rm ad} = 1k_BT$, $F = 3k_BT\ l_{\rm min}^{-1}$ and $\rho = 3.45\%$. Three panels for three different shear conditions. Without shear, the vesicle retains its position under thermal fluctuation. In presence of shear, it moves with the flow.

**Movie 3: Non-motile vesicle** Non-motile weakly spreading vesicle is obtained $E_{\rm ad} = 1k_BT$, $F = 0.5k_BT\ l_{\rm min}^{-1}$ and $\rho = 3.45\%$. Three panels for three different shear conditions. The vesicle moves with the shear flow, however, it remains at the same position without shear flow because the vesicle is non-motile.

**Movie 4: Pancake vesicle** Non-motile pancake-shaped vesicle is obtained $E_{\rm ad} = 0.5k_BT$, $F = 2k_BT\ l_{\rm min}^{-1}$ and $\rho = 6.91\%$. Three panels for three different shear conditions. The vesicle slides with the shear flow whereas no shear condition leads the vesicle to remain at the initial position without moving significantly.

---

[1] I. Cantat and C. Misbah. Lift force and dynamical unbinding of adhering vesicles under shear flow. *Phys. Rev. Lett.*, 83:880–883, Jul 1999.

[2] M. Fošnarič, S. Penič, A. Iglič, V. Kralj-Iglič, M. Drab, and N. S. Gov. Theoretical study of vesicle shapes driven by coupling curved proteins and active cytoskeletal forces. *Soft Matter*, 15(26):5319–5330, 2019.

[3] R. K. Sadhu, S. Penič, A. Iglič, and N. S. Gov. Modelling cellular spreading and emergence of motility in the presence of curved membrane proteins and active cytoskeleton forces. *The European Physical Journal Plus*, 136(5):495, 2021.
